# Long-term exposure to PFE-360 in the AAV-α-synuclein rat model: findings and implications

**DOI:** 10.1101/465567

**Authors:** Michael Aagaard Andersen, Florence Sotty, Poul Henning Jensen, Lassina Badolo, Ross Jeggo, Garrick Paul Smith, Kenneth Vielsted Christensen

## Abstract

Treatment of Parkinson’s disease is reliant on symptomatic treatments, without any option to slow or halt disease progression. Mutations in LRRK2 and α-synuclein are known risk factors for Parkinson’s disease. Presence of α-synuclein aggregates at autopsies in both idiopathic and most G2019S cases is suggestive of a common disease pathogenesis. LRRK2 and α-synuclein interaction is hypothesized to play a pivotal role in the pathological mechanisms and inhibitors of LRRK2 are investigated as novel disease modulatory treatments in the clinic. However, preclinical *in vivo* evidence of a beneficial effect of LRRK2 inhibition is mixed and limited. This study increases our understanding of LRRK2 as a mediator of neuronal dysfunction and the potential of LRRK2 as a promising target in PD.

## Introduction

Historically, neurodegeneration of dopaminergic neurons in substantia nigra pars compacta and α-synuclein inclusions in Lewy bodies and Lewy neurites are the histopathological hallmarks of Parkinson’s disease (PD) (Spillantini et al., 1997; Antony et al., 2013). According to the hypothesis raised by Braak and Beach, pathogenic α-synuclein species may be capable of transmitting their pathological properties across brain nuclei following neuronal/axonal pathways (Braak et al., 2003, 2004; Beach et al., 2009). A number of known dominant autosomal missense mutations, duplications and triplications in the SNCA gene encoding α-synuclein are risk factors for developing PD (Pankratz et al., 2007; Lill, 2016). The exact physiological role of α-synuclein is not fully understood. However, virally and genetically manipulated animal models have revealed impairments in synaptic transmission and vesicular machinery (Cabin et al., 2002; Yavich et al., 2004; Watson et al., 2009; Nemani et al., 2010; Busch et al., 2014; Subramaniam et al., 2014), suggesting an involvement of α-synuclein in normal synaptic neurotransmission.

Increased leucine-rich repeat kinase 2 (LRRK2) kinase activity is suggested as the key risk factor associated with late onset PD (Funayama et al., 2002; Zimprich et al., 2004b; Paisán-Ruíz et al., 2004; Zimprich et al., 2004a; Di Fonzo et al., 2005, 2006; Gilks et al., 2005; Kachergus et al., 2005; Nichols et al., 2005; Healy et al., 2008; Ross et al., 2011). The physiological function of LRRK2 kinase has recently been assigned as a controller of RAB GTPase activity via increased phosphorylation of several small RAB GTPase, and autophosphorylation (LRRK2-pS1292) of disease relevant mutant LRRK2 (Steger et al., 2016, 2017; Fan et al., 2017; Liu et al., 2017; Thirstrup et al., 2017). Genetic ablation and pharmacological inhibition of LRRK2 have previously been demonstrated to have a promising effect on disease relevant alterations in preclinical *in vivo* models of PD (Lin et al., 2009; Daher et al., 2014, 2015; Andersen et al., 2018a). The partly shared clinical and histopathological manifestation of idiopathic/sporadic and LRRK2 PD is suggestive of at least some shared physiopathological mechanisms between LRRK2 PD and sporadic PD, thereby supporting a potential for LRRK2 inhibition in the modulation of common pathways in the disease mechanisms.

Glutamatergic neurons in the STN are a key regulator of neuronal input to the motor thalamus (Nambu et al., 2002). In PD patients and neurotoxin as well as viral models of PD, the loss of striatal dopaminergic (DAergic) neurotransmission triggers an increase in STN burst discharge pattern, and aberrant oscillations in the beta range throughout the basal ganglia (Brown, 2007; Steigerwald et al., 2008; Wilson et al., 2011; McConnell et al., 2012; Pan et al., 2016; Andersen et al., 2018a). Functional pre-clinical studies and deep brain stimulation studies in both PD models and humans suggest that functional relief of aberrant STN burst firing is associated with acute normalization of motor function (Grill et al., 2004; Benabid et al., 2009; McConnell et al., 2012; Moran et al., 2012; Pan et al., 2016; Wichmann and DeLong, 2016). Recently, acute effects of LRRK2 inhibition have been reported on α-synuclein-induced aberrant STN burst firing (Andersen et al., 2018a), possibly representing a disease relevant measurement to evaluate new disease modulatory targets.

In the present study, AAV-α-synuclein (human wildtype) overexpression in rat substantia nigra recapitulates PD-like aberrant STN burst firing, DAergic neurodegeneration and motor dysfunction. Using the AAV-α-synuclein overexpression approach, we investigated the effect of chronic LRRK2 inhibition on α-synuclein-induced neuronal dysfunction. Chronic LRRK2 inhibition had a partial restorative effect on motor function, which was not correlated with any effect on aberrant STN firing, dopaminergic neurodegeneration measured by striatal tyrosine hydroxylase expression, LRRK2 expression, α-synuclein expression or phosphorylation (α-synuclein-pS129). The present findings do not strongly support the mechanistic link between LRRK2 kinase activity and pathological α-synuclein, and further reveal some limitations of the preclinical AAV α-synuclein overexpression model.

## Methods

### In vivo experimental methods

#### Ethical statement

All experiments were carried out in accordance with the European Communities Council Directive (86/609/EEC) for the care and use of laboratory animals and the Danish legislation regulating animal experiments.

#### Animals

Female Sprague Dawley rats were acquired from Taconic Denmark (NTac:SD, n=23 in total). Rats were group housed in humidity (55-65 %) and temperature (22 ± 1.5 °C) controlled conditions in a 12:12 hours light/dark cycle. Water and food were available *ad libitum*. Female rats were used as experimental subject as they have lower body weight and lower variation in skull size compared to male rats at 26 weeks of age where the experiment was carried out. Lower body weight and variation in skull size are preferred in the *in vivo* electrophysiological experiments.

#### Viral mediated α-synuclein overexpression in substantia nigra

Injection of recombinant adeno-associated-viral vectors (AAV) was performed in 10-12 weeks old animals (weighing 250-300 g). Surgical neuroleptic analgesia was induced with a mixture of Hypnorm^®^ and Dormicum (corresponding to 157 µg/kg fentanyl, 2.5 mg/kg midazolam and 5 mg/kg fluanisone). All incisions were infiltrated with local analgesia (lidocaine). The rats were placed in a stereotaxic frame and the body temperature was maintained at 37.5 °C. After a skin incision, a hole was drilled in os paritalis directly above the substantia nigra pars compacta (SNc) at the following coordinates, according to Paxinos and Watson (2007): 5.5 mm caudal to bregma and 2.0 mm lateral to the midline. The injection cannula (Hamilton, 30 gauge stainless steel cannula) was slowly lowered into the injection site located 7.2 mm ventral to the brain surface. A unilateral injection of 3 µl AAV viral vector (3 x 10^10^ GC/ml) (Vector Biolabs, Malvern, PA, USA) containing the transgene for human wildtype (hwt) α-synuclein or an empty vector (CTRL) was performed at a flow rate of 0.2 µl/min. The cannula was left in place for 5 min after the injection to allow diffusion. After the skin was sutured, a 24 hours analgesic treatment protocol was initiated (buprenorphine 0.05 mg/kg every 8^th^ hour). The transgene expression was driven by a chimeric promoter element containing part of the Cytomegalovirus promoter and a part of the synthetic chicken β-actin promoter. Further, the expression was enhanced by the woodchuck of hepatitis virus post-translational regulatory element. A successful viral injection was further validated by detection of hwt α-synuclein in the ipsilateral striatum by SDS-page and subsequent Western blot.

#### PFE-360 treatment

PFE-360 (Baptista et al., 2015) was synthesized at H. Lundbeck A/S as described in patent application US2014/0005183 (Example 217) (1-methyl-4-(4-morpholino-7H-pyrrolo[2,3-d] pyrimidin-5yl) pyrrole-2-carbonitrile). PFE-360 was dissolved at a concentration of 3 mg/ml in a vehicle containing 10 % captisol and adjusted to pH 2 with 1 M methane sulfonic acid. The solution was administrated by oral gavage using plastic feeding tubes. Rats receiving PFE-360 were dosed with 7.5 mg/kg twice daily with a 12 hours interval. The treatment was initiated on day 1 post-AAV injection. The dose of 7.5 mg/kg was chosen based on a separated pharmacokinetic study showing that 7.5 mg/kg gives full LRRK2 kinase inhibition from 1-10 hours and approximately 50 % LRRK2 kinase inhibition at 12 hours post dosing. The plasma pharmacokinetic profile of PFE-360 can be found elsewhere (Andersen et al., 2018b).

#### Motor symmetry assessment in the cylinder test

Nine weeks after the viral injection, motor function was assessed in the cylinder test. Rats receiving PFE-360 or vehicle were tested 1-2 hours post dosing. Briefly, the rat was placed in a transparent cylinder and video recorded for 5 min. A minimum of 15 touches was required to ensure the sensitivity of the test. The ratio between the contralateral forepaw (to the injection) and the total number of touches was used as the primary read out.

#### Spontaneous locomotor activity

Following the cylinder test, the non-forced spontaneous locomotor activity was evaluated in a cage similar to the home cage with clean bedding but without enrichment for 1 hour. Activity was recorded by 4 photosensors along the cage. There was no habituation period. The activity count was calculated as the number of beam breaks, excluding repeated break of the same beam. Rats receiving PFE-360 or vehicle were tested 5-7 hours post-dosing.

#### Single unit recordings of glutamatergic subthalamic neurons

10-12 weeks following the viral vector injection, the same animals were subjected to single unit recording of putative glutamatergic STN neurons under urethane anesthesia (1.6 – 1.9 g/kg, i.p.). The level of anesthesia was maintained and monitored as absence of the deep pain reflexes but presence of the corneal reflex. This allowed to ensure that recordings were performed under similar levels of anesthesia, as the relative brain state influences the basal ganglia firing properties (Magill et al., 2000). The rat was placed in a stereotaxic frame and the body temperature was maintained at 37.5 ± 0.5 °C. After a midline incision, a hole was drilled in os paritalis above the STN ipsilateral to the viral injection, at the following coordinates: 3.0 - 4.2 mm caudal to bregma and 2.4 – 2.8 mm lateral to the midline (Paxinos and Watson, 2007). The brain surface was kept moist with 0.9% saline after removal of the dura mater. The recording electrodes were fabricated from borosilicate glass capillaries (1B150F-4, World Precision Instruments) pulled into a fine tip under heating and broken under a microscope to a tip diameter of 3-8 µm reaching an *in vitro* resistance of 3 – 9 MΩ. The electrode was filled with 0.5 M sodium acetate containing 2% of Pontamine Sky Blue. The electrode was slowly lowered into STN using a motorized micromanipulator. STN was typically localized at 6.6 – 7.8 mm ventral to the brain surface (Paxinos and Watson, 2007). The action potentials were amplified (x 10k), band pass filtered (300Hz – 5kHz), discriminated and monitored on an oscilloscope and an audio monitor. Spike trains of action potentials were captured and analyzed using Spike 2 v 7.13 software with a computer-based system connected to the CED 1401 (Cambridge Electronic Design Ltd., Cambridge, UK). A minimum of 500 consecutive spikes was used for analysis of basal firing properties. PFE-360 was administrated to the rats per oral 1 hour before the start of the recording session and ended 6 – 8 hours post-dosing.

At end of the recording session, an iontophoretic ejection of Pontamine Sky Blue was achieved by applying a negative voltage to the electrode allowing visualization of the last recording position using classical histological methods. The recording sites were then retrospectively reconstructed, and neurons were only included in the analysis if the reconstruction provided evidence of the neuron to be within the STN.

#### Single unit recordings data analysis

The action potential of glutamatergic neurons in STN is typically short (<2 ms) with a biphasic waveform (Hollerman and Grace, 1992). Each spike-train was carefully inspected before the analysis using the principle component tools and the overdraw wavemark function in Spike 2 v 7.13, making sure that only spikes from a single neuron were included in the analysis. The signal to noise ratio was always >2:1. Spike trains were analyzed offline using a custom-made script measuring the firing rate and coefficient of variation of the interspike interval (CV ISI). Briefly the CV ISI is defined as the ratio between the standard deviation of the interspike interval and the average interspike interval x 100 (Herrik et al., 2010). Further, the classification of the neuronal firing pattern was divided into regular, irregular and bursty based on a visual inspection of the autocorrelogram and the discharge density histogram (Tepper et al., 1995; Kaneoke and Vitek, 1996).

### Ex-vivo biochemical methods

#### Brain sampling

At the end of the recording session, the rats were perfused transcardially with 100 ml 0.9 % saline containing 0.3 % heparin. The brain was removed and sectioned on an ice-cold plate. The striata were snap frozen on dry-ice and stored at – 80 °C until preparation of tissue lysates for SDS-PAGE and Western blot. The caudal part of the brain containing STN was stored at – 20 °C for histological validation of the recording sites. The cerebellum was weighted and snap frozen in a Covaris tube on dry ice and stored at – 80 °C until further analysis.

#### SDS-PAGE and Western blot sample preparation

Preparation of striatal samples for SDS-page and Western blot was performed as described elsewhere (Andersen et al., 2018a).

#### SDS-page and Western blot quantification of tyrosine hydroxylase, striatal enriched phosphatase, human wild-type α-synuclein and phosphorylated α-synuclein-S129

Quantification of TH, STEP and hwt-α-synuclein expression and α-synuclein-S129 phosphorylation were done as described elsewhere ((Andersen et al., 2018a). Briefly, the quantification was done using SDS-page and subsequent Western blot. Finally, the proteins were visualized quantified by infrared detection using Li-Cor Odyssey CLx (Li-COR, Nebraska, US).

#### Total LRRK2 and phosphorylated LRRK2-S935 quantification

Total LRRK2 expression levels and the level of LRRK2 inhibition were quantified using SDS-page and subsequent Western blot as described elsewhere (Andersen et al., 2018b). The total LRRK2 was quantified for both ipsi- and contralateral striatum and reported as the mean of the two measurements.

#### Quantification of PFE-360 brain exposure

The analysis method is described elsewhere (Andersen et al., 2018a). Briefly, the quantification of PFE-360 brain exposure was performed using a LC-MS/MS coupled to a Waters Acquity UPLC.

### Statistical analysis

Neuronal firing properties (firing rate and CV ISI) data were analyzed using Kruskal-Wallis test with Dunn’s corrected p-value for multiple comparison, as they were non-normally distributed. Distributive differences in firing pattern were statistically assessed using Chi-squared analysis. The p-value for multiple chi-squared tests were corrected using the Bonferroni correction (p > 0.0125 was considered significant). Motor behavior and Western blot quantifications (TH, STEP, α-synuclein-pS129, LRRK2 and LRRK2-pS935) data were all normally distributed and were analyzed using a one-way ANOVA test with Tukey’s post hoc multiple comparison. Statistical analysis of human α-synuclein overexpression and unbound PFE-360 exposure were done using an unpaired t-test. All statistics and figures were made in GraphPad Prism^®^ 7.03. All values are presented as mean ± SEM.

## Results

### Chronic PFE-360 dosing modulates AAV-α-synuclein-induced motor deficit

In our effort to evaluate the therapeutic preclinical potential of chronic LRRK2 inhibition, we first tested the effect of PFE-360 (7.5 mg/kg BID) on motor symmetry in the cylinder test. Overall, there was a statistically significant effect on the forepaws ratio across groups (Fig. 1A). Multiple comparisons showed that AAV-overexpression of α-synuclein induced a significant asymmetry in the motor function compared to an empty vector in vehicle-treated animals. Chronic dosing with PFE-360 did not affect the motor symmetry assessed in the cylinder test in animals injected with an empty vector. In rats injected with AAV-α-synuclein, PFE-360 partially normalized the dysfunctional use of forepaws as evidenced by the significantly increased the use of the contralateral forepaw compared to vehicle treatment.

**Figure 1.**
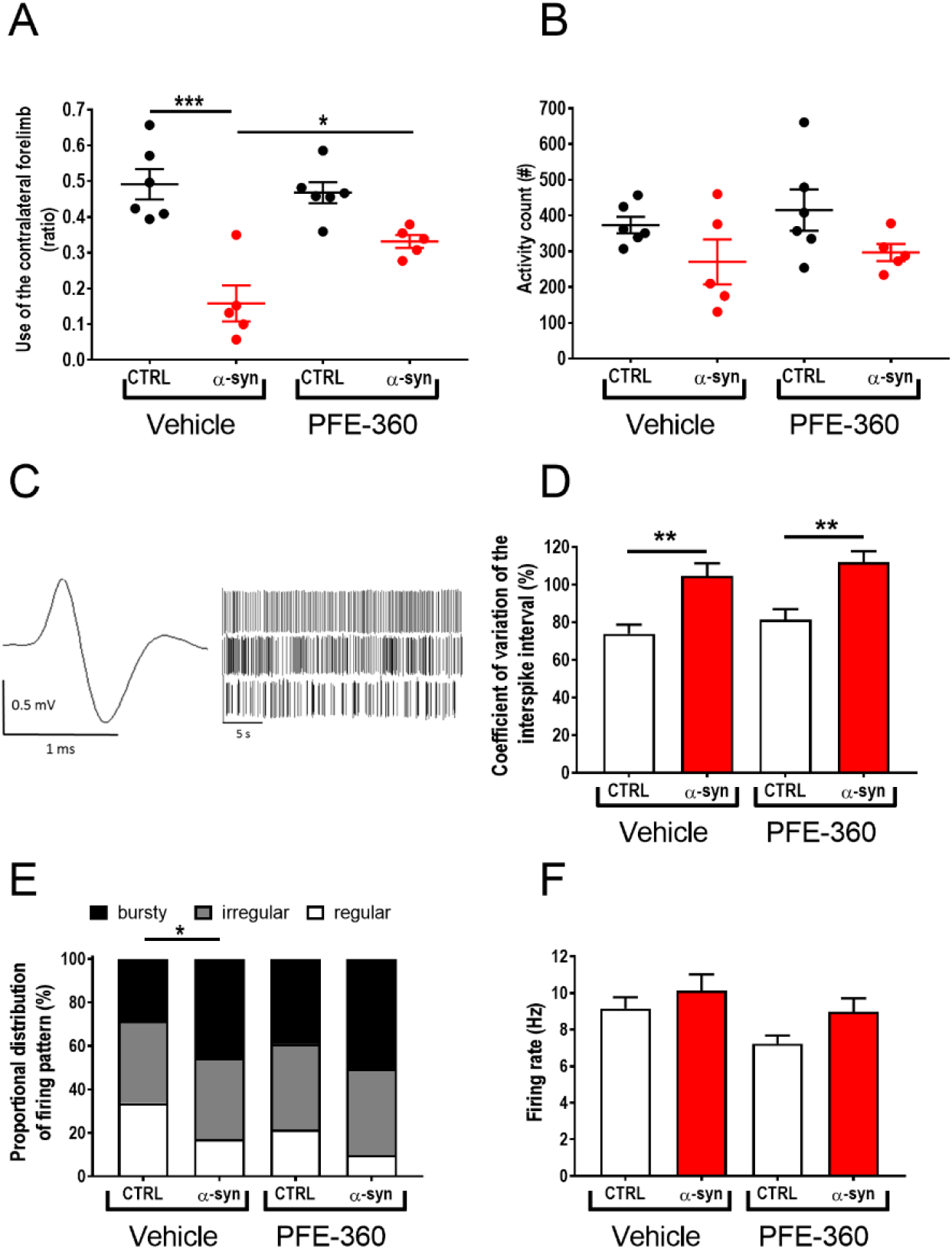
Chronic PFE-360 treatment partially restores motor function independently of STN burst firing induced by AAV-α-synuclein overexpression. **(A)** Motor symmetry assessed 9 weeks post intra-cerebral viral inoculation of AAV-α-synuclein or CTRL (empty AAV-vector). The test was performed 1-2 hours post-dosing (one-way ANOVA, p < 0.001) (CTRL veh and PFE-360 N = 6, α-syn veh and PFE-360 N = 5). **(B)** Locomotor activity measured 5-7 hours post-dosing revealed a non-significant trend for a decreased activity in α-synuclein overexpressing rats in both treatment groups compared to CTRL rats. PFE-360 treatment did not have any adverse effect on locomotion in the CTRL group (one-way ANOVA, p = 0.51) (CTRL veh and PFE-360 N = 6, α-syn veh and PFE-360 N = 5). **(C)** Representative action potential and spike-trains from single unit recordings of putative glutamatergic neuron in the STN. The spike-trains are representative examples of regular (top), irregular (middle) and bursty (bottom) firing patterns. **(D)** Coefficient of variation of the interspike interval is increased by α-synuclein overexpression in both treatment groups (Kruskal-Wallis test, p < 0.001). **(E)** Proportional distribution of regular, irregular and bursty neurons (χ^2^ = 22.28, Df = 6, p = 0.0011) shows increased proportion of bursty neurons in α-synuclein overexpressing rats in both treatment groups. **(F)** The average firing rate of STN neurons is not significantly different between groups (Kruskal-Wallis test, p =0.11). DF: (N: number of animals; n: number of neurons; CTRL veh (N = 5, n = 98), α-synuclein veh (N = 6, n = 99), CTRL PFE-360 (N = 6, n = 90) and α-synuclein PFE-360 (N = 6, n = 113). Tukey’s multiple comparison is represented by the lines. *p < 0.05, **p < 0.01 and ***p < 0.001. For multiple Chi-squared tests in (E) the Bonferroni-corrected p-value was used, *p < 0.0125 (4 comparisons: significance level p = 0.0125).

Analysis of the spontaneous locomotor activity 5-7 hours after the last dosing did not show any overall statistical effect of either the viral vector or the treatment, although a trend for reduced locomotor activity was observed in animals injected with AAV-α-synuclein compared to an empty vector (Fig 1B).

### AAV-α-synuclein-induced aberrant STN burst firing is not attenuated by chronic LRRK2 inhibition using PFE-360

To investigate the modulatory effect of repeated LRRK2 inhibition on STN burst firing induced by hwt α-synuclein overexpression in substantia nigra, rats were repeatedly dosed with PFE-360 for 10-12 weeks following viral injection. The electrophysiological properties of putative glutamatergic neurons in STN were recorded at the end of the treatment period, and 1-8 hours following the last dose. A total of 400 putative glutamatergic STN neurons from 22 rats divided into 4 groups were included in the final analysis (Fig. 1C-F). Overall, comparison of the STN firing pattern revealed a significant group effect in the distribution of regular, irregular and bursty firing neurons between groups and a significant group effect in the CV ISI (Fig. 1D + E). Multiple comparisons revealed that overexpression of α-synuclein significantly increased the proportion of burst firing neurons and the CV ISI compared to an empty vector in vehicle-treated animals. In empty vector-injected animals, chronic dosing with PFE-360 7.5 mg/kg BID did not alter the relative distribution of the firing pattern or the CV ISI (CTRL vehicle vs. CTRL PFE-360 7.5 mg/kg). In AAV-α-synuclein injected animals, chronic dosing with PFE-360 did not have any significant effect on the relative distribution of the firing pattern or the CV ISI. The CV ISI was significantly increased compared to CTRL PFE-360, to a level comparable to α-syn PFE-360. The relative distribution of firing pattern of α-synuclein was not significantly altered compared to CTRL PFE-360 or α-synuclein vehicle groups. The firing rate was not overall significant different between groups (Fig. 1F).

### Striatal LRRK2 expression is not decreased following chronic LRRK2 inhibition

Evaluation of the impact of PFE-360 on total LRRK2 expression levels in striatum revealed an overall significant difference between groups (Fig. 2A + B). Post hoc analysis revealed no significant effect of AAV-α-synuclein overexpression in vehicle treated rats, nor was there any significant effect of PFE-360 treatment in CTRL rats. However, PFE-360 treatment in AAV-α-synuclein overexpressing rats significantly decreased the LRRK2 expression level compared to vehicle treated α-synuclein overexpressing rats. There was no statistical difference between PFE-360 treated CTRL and α-synuclein rats.

**Figure 2.**
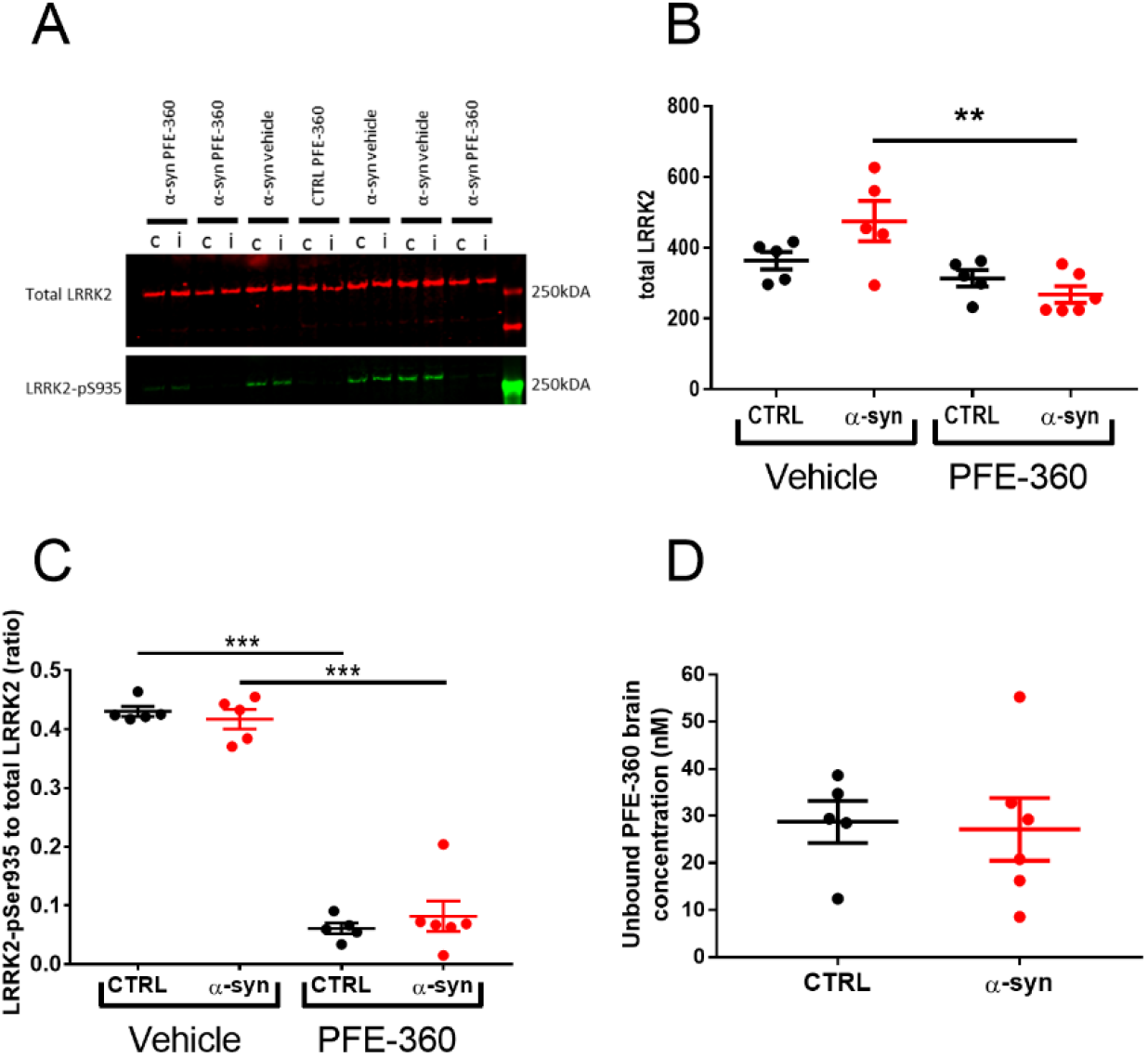
Striatal LRRK2 expression is not changed by PFE-360 treatment despite full LRRK2 inhibition. **(A)** Western blot analysis of total LRRK2 and phosphorylated LRRK2-S935 (LRRK2-pS935) after 10-12 weeks chronic PFE-360 dosing. The tissues were collected 6-8 hours after last dosing. i = ipsilateral to the viral injection and c = contralateral to the viral injection. **(B)** Quantification of Western blots shows that chronic PFE-360 treatment did not affect LRRK2 expression in CTRL rats compared to vehicle treatment. In α-syn rats, PFE-360 treatment significantly lowered LRRK2 expression compared to vehicle treatment, although each α-syn group did not differ significantly from their respective CTRL group (one-way ANOVA, p = 0.0032). **(C)** The LRRK2-pS935 to total LRRK2 ratio was used as a measure of target engagement. At full inhibition, the ratio is typically <0.1 and was achieved in both CTRL and α-syn rats treated with PFE-360 (one-way ANOVA, p <0.001). **(D)** Unbound PFE-360 brain concentrations 6-8 hours post-dosing. IC_50_ of PFE-360 was previously calculated to 2.3 nM giving theoretical 99 % LRRK2 inhibition at 23 nM (unpaired t-test, p = 0.85). CTRL veh, CTRL PFE-360 and α-syn veh; N = 5, α-syn PFE-360 N = 6. Tukey’s multiple comparison is represented by the lines. **p < 0.01 and ***p < 0.001.

### PFE-360 exposure profile and target occupancy in rat brain

LRRK2 engagement and kinase inhibition in striatum of PFE-360 6-8 hours post-dosing was evaluated using SDS-page and Western blotting (Dzamko et al., 2010, 2012; Nichols et al., 2010). Viral α-synuclein overexpression did not impact levels of LRRK2-pS935 or total LRRK2 ratio (Fig. 2C). In CTRL and α-synuclein rats dosed with PFE-360 significant decreases in LRRK2-pS935 to total LRRK2 ratio were observed. The result suggests that extended dosing of PFE-360 at 7.5 mg/kg for 10-12 weeks gives full LRRK2 kinase inhibition 6-8 hours post-dosing. A single rat exhibiting a PFE-360 exposure level approximately 3 x IC_50_, suggestive of 70% inhibition at 8 hours post-dosing, was nevertheless included in the study (Fig. 2D). The calculated LRRK2 IC_50_ value and inhibition level is based on a single dose study using 7.5 mg/kg PFE-360 (Andersen et al., 2018a). Based on the exposure levels following a single PFE-360 dose of 7.5 mg/kg, the average level of LRRK2 inhibition in the current study is estimated to be from 100 % to 50 % over 24 hours, fluctuating between 100 % at 1-10 hours post-dosing and 100-50 % at 10-12 hours post-dosing (data shown elsewhere (Andersen et al., 2018a)).

### LRRK2 inhibition does not attenuate the loss of TH expression in striatum induced by α-synuclein overexpression

Neurodegenerative processes induced by AAV-α-synuclein overexpression were assessed indirectly by quantifying the expression level of TH in striatum. Overall, there was a significant effect of groups on striatal TH expression levels (Fig. 3A + B). Post hoc analysis revealed a significant loss of striatal TH induced by AAV-α-synuclein overexpression compared to an empty vector independently of treatment.

**Figure 3.**
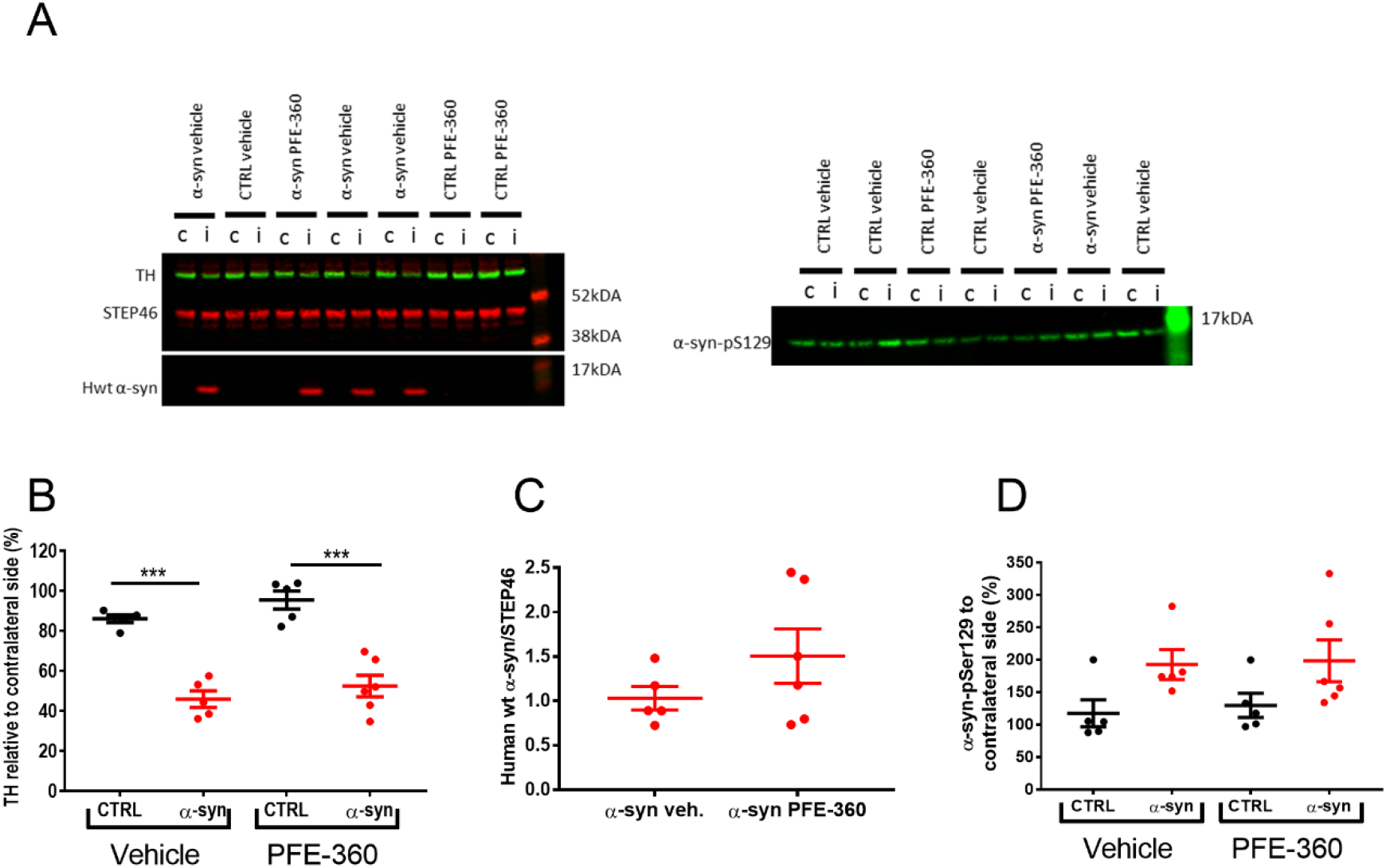
Striatal neurodegenerative processes and α-synuclein phosphorylation are not halted by chronic LRRK2 inhibition. **(A)** Western blot analysis of tyrosine hydroxylase (TH) and human wild-type α-synuclein (hwt α-syn) (left) as well as phosphorylation of α-synuclein-S129 (α-syn pS129) (right). i = ipsilateral to the viral injection and c = contralateral to the viral injection. **(B)** Quantification of TH expression from ipsi- and contralateral striatum normalized to STEP46 and presented as percentage of contralateral striatum shows a significant loss of TH expression in α-syn overexpressing rats in both treatment groups (one-way ANOVA, p < 0.001). **(C)** The expression level of hwt α-synuclein normalized to STEP46 is not significantly different between groups (unpaired t-test, p = 0.22). **(D)** The α-synuclein-S129 phosphorylation from ipsi- and contralateral striatum normalized to STEP46 and presented as percentage of contralateral striatum is not significantly different between groups (one-way ANOVA, p = 0.076). CTRL veh, CTRL PFE-360 and α-syn veh; N = 5, α-syn PFE-360 N = 6. Tukey’s multiple comparison is represented by the lines. ***p < 0.001.

### LRRK2 inhibition does not affect human wt α-synuclein expression or phosphorylation of total α-synuclein

The overexpression of hwt α-synuclein was not significantly different between PFE-360 and vehicle-treated animals (Fig. 3A + C). The levels of pS129-α-synuclein in the striatum were overall not significantly different across groups (Fig 3A + D). A trend for increased expression was observed following AAV-α-synuclein overexpression compared to CTRL animals although this was not significant. Chronic LRRK2 inhibition with PFE-360 did not impact α-synuclein-pS129 levels compared to vehicle in CTRL or AAV-α-synuclein rats.

## Discussion

Firstly, the findings suggest that the use of our current preclinical PD model in terms of predicting LRRK2 inhibitor efficacy in the clinic should be highly cautioned. Secondly, further investigation into mechanisms underlying these differences between models and investigators is needed before any solid conclusions can be made on the translational value of our current preclinical PD model. Finally, the discrepancy between the reported acute and present chronic effects of LRRK2 inhibition on aberrant α-synuclein-induced STN firing activity represents an interesting aspect of LRRK2 as a regulator of basal ganglia neurotransmission. Previous reports using acute or chronic LRRK2 inhibition have shown preclinical modulatory and protective effects of LRRK2 inhibition in α-synuclein-based PD models, supporting a pathophysiological interaction between LRRK2 kinase function and α-synuclein-induced pathology (Lin et al., 2009; Daher et al., 2015; Andersen et al., 2018a). In the present report, chronic LRRK2 inhibition had a restorative effect on the motor performance evaluated in the cylinder test. Interestingly, the PFE-360-induced improvement in motor function was not correlated to a similar reversal of STN burst firing. This suggests that chronic LRRK2 inhibition may exert effects on motor function through action on other pathways bypassing STN, e.g. affecting neurotransmission and/or synaptic plasticity within the motor circuit, thereby functionally circumventing the relative strength/importance of STN firing on the motor output. The complexity of the motor circuit, where timing of a signal through the four pathways in the basal ganglia are equally important in determining the motor outcome (Nambu et al., 2002; Mastro and Gittis, 2015), may contribute to the discrepancy between the effects of PFE-360 on motor behaviour and STN activity. Changes in synaptic plasticity have been reported in transgenic animals overexpressing LRRK2 (Beccano-Kelly et al., 2015; Sweet et al., 2015), but no data from LRRK2 knock out animals is available. In the latter, it is conceivable that LRRK2 ablation from conception may induce compensatory mechanisms which would not be observed in wild-type animals unless long-term pharmacological inhibition would trigger similar compensations. Another possibility is that the function of LRRK2 inhibition needs an external trigger to reveal its functional involvement in neurotransmission such as impaired DAergic neurotransmission in substantia nigra and striatum. In this respect, alterations in glutamatergic transmission in striatal slices from G2019S-LRRK2 transgenic animals were only observed when rotenone, a mitochondrial toxin, was also present, indicating that LRRK2 might only play a role in the diseased state, such as impaired DAergic neurotransmission (Tozzi et al., 2018). In support, recent studies have suggested that LRRK2 kinase activity is increased in substantia nigra in the AAV-α-synuclein overexpression model and in rotenone models as well as in postmortem tissue from idiopathic PD patients (Howlett et al., 2017; Di Maio et al., 2018).

The acute LRRK2 inhibition with PFE-360 was shown to drastically reduce the spontaneous locomotor activity of control animals (Andersen et al., 2018a), a hypo-locomotor effect was not observed after repeated PFE-360 treatment in the present study. This potentially indicates an adaptation to the hypolocomotor effect after repeated administration. Our target engagement measurements showed full LRRK2 inhibition in the brain after repeated treatment (and periphery (data not shown), see (Andersen et al., 2018b)), but no consistent change in LRRK2 expression, thereby ruling out adaptive effects mediated directly by LRRK2 and further suggesting the existence of adaptive mechanisms taking place downstream of LRRK2. Further investigation in long term treated subjects preclinically and clinically will be needed to validate such physiological effects.

Our investigation recapitulated the effect of AAV-α-synuclein overexpression on STN basal firing properties by increasing the proportion of burst firing neurons, mimicking findings in neurotoxin models and PD patients (Hollerman and Grace, 1992; Bergman et al., 1994; Ni et al., 2000; Iancu et al., 2005; Brazhnik et al., 2014; Pan et al., 2016). As above mentioned, chronic LRRK2 inhibition using PFE-360 was unable to counteract AAV-α-synuclein-induced aberrant STN burst firing. Previously, aberrant STN firing in the AAV-α-synuclein rat model is reported to be normalized by acute LRRK2 inhibition using PFE-360 (Andersen et al., 2018a). In the present study, all putative STN neurons were recorded under 70 – 100% LRRK2 inhibition (based on theoretical back calculations of brain exposure over time (data not shown)). The discrepancy between the effect of acute and chronic LRRK2 inhibition on STN firing properties further supports induction of downstream adaptation to PFE-360 after long term exposure, which was also observed in the locomotor test.

We further showed that chronic treatment with PFE-360 did not have any impact on DAergic neurodegeneration, striatal hwt-α-synuclein expression or striatal α-synuclein-pS129 level induced by AAV-α-synuclein overexpression in substantia nigra and associated pathways. These findings contrast findings by Daher et al who reported protection of DAergic neurons from α-synuclein-induced neurodegeneration after chronic treatment with a similar LRRK2 inhibitor chemotype (Daher et al., 2015). A possible explanation for this discrepancy is related to the different design of the respective studies. In the study by Daher et al. (2015), animals were assessed 4 weeks following viral injection and drug treatment initiation, which may have allowed revealing a delay in the progression of the pathology. The viral serotype used was also different in their study compared to the present, which can have important impact on the infectious potential of the viral vector (Van der Perren et al., 2011). Importantly, the treatment regimen used in our study did not result in full LRRK2 inhibition throughout the whole duration of the treatment based on the free brain concentration of PFE-360 at 12 hours post-dose (data from acute administration (Andersen et al., 2018a)). Whether full brain LRRK2 inhibition was achieved throughout the study reported by Daher et al. (2015) is unknown. Thus, to fully explore the potential of LRRK2 kinase inhibitors in preclinical models, further development of novel LRRK2 inhibitors with more optimal pharmacokinetic and pharmacodynamic properties is needed or a study design reaching full LRRK2 inhibition 24 hours a day.

Chronic treatment with PFE-360 in CTRL rats was not associated with a change in LRRK2 expression in the striatum *in vivo*. The statistical effect on total LRRK2 expression was seen between groups overexpressing α-synuclein treated with PFE-360 or vehicle. Previous studies in the AAV-α-synuclein rat model have not found similar effects of α-synuclein overexpression (Andersen et al., 2018a). The lack of changes in LRRK2 expression after chronic LRRK2 inhibition in CTRL rats is in agreement with similar findings in the brain after 11 days chronic treatment with MLi-2 (Fell et al., 2015). In contrast, others have reported large decreases in total brain LRRK2 expression *in vivo* and *in vitro* in cellular systems after LRRK2 kinase inhibition (Herzig et al., 2011; Daher et al., 2015; Fuji et al., 2015; Zhao et al., 2015; Lobbestael et al., 2016). In this regard, the species (rat vs mouse vs non-human primate) and LRRK2 expression level should be considered. *In vitro* overexpression of LRRK2 might be very different from *in vivo* conditions with much lower expression of LRRK2 *in vivo* and might represent different mechanisms not allowing direct comparison.

Although the viral overexpression model has considerably increased our understanding of α-synuclein-induced alterations in the basal ganglia circuitry, the α-synuclein preformed fibrils model has been recently suggested to represent a more faithful model of idiopathic PD (Duffy et al., 2018), and may therefore be highly valuable to further validate LRRK2 as a therapeutic target for idiopathic PD.

## Summary

We have demonstrated that chronic LRRK2 inhibition induced a partial reversal of the motor dysfunction in the AAV-α-synuclein rat model of PD. However, this effect was not associated with any beneficial effect on STN burst firing or loss of striatal TH expression. The discrepancies between previously reported acute effects and the present chronic effects of LRRK2 inhibition are likely to be dependent on compensatory mechanisms after chronic LRRK2 inhibition. Nevertheless, the improvement in motor function remains an interesting finding. In this respect, further studies in other preclinical models such as the α-synuclein PFF model may prove valuable.

In the search for the precise mechanisms underlying a putative interaction between LRRK2 and α-synuclein, changes in synaptic plasticity and vesicular trafficking seems to play a pivotal role in model systems manipulating neuronal and non-neuronal LRRK2 kinase function, respectively. Such changes might result in wanted or unwanted effects. It was previously described that LRRK2 is involved in synaptic plasticity and that it is dysregulated in the parkinsonian state (Beccano-Kelly et al., 2015; Chu et al., 2015; Mastro and Gittis, 2015; Sweet et al., 2015) thereby strengthening a potential beneficial effect of LRRK2 inhibition on synaptic plasticity in the pathological conditions.

## Acknowledgements

The authors acknowledge Camilla Thormod Haugaard and Bo Albrechtslund for their expertise and technical assistance. Financial support was provided by the Danish Ministry of Science and Innovation (ID: 16448). The authors declare no competing financial interests.

